# Focal Deletions of a Promoter Tether Activate the *IRX3* Oncogene in T Cell Acute Lymphoblastic Leukemia

**DOI:** 10.1101/2024.02.06.579027

**Authors:** Sunniyat Rahman, Gianna Bloye, Nadine Farah, Jonas Demeulemeester, Joana R. Costa, David O’Connor, Rachael Pocock, Adam Turna, Lingyi Wang, SooWah Lee, Adele K. Fielding, Juliette Roels, Roman Jaksik, Malgorzata Dawidowska, Pieter Van Vlierberghe, Suzana Hadjur, Jim R. Hughes, James O.J. Davies, Alejandro Gutierrez, Michelle A Kelliher, Peter Van Loo, Mark A. Dawson, Marc R. Mansour

**Affiliations:** University College London Cancer Institute, Department of Haematology, London, UK; Peter MacCallum Cancer Centre, Melbourne, Victoria 3000, Australia; Sir Peter MacCallum Department of Oncology, The University of Melbourne, Victoria 3010, Australia; VIB KU Leuven Centre for Cancer Biology, 3000 Leuven, Belgium; Department of Developmental Biology and Cancer, GOS Institute of Child Health, UCL; University of York, Hull York Medical School, UK; Ghent University, Department of Biomolecular Medicine, Ghent, Belgium; Department of Systems Biology and Engineering and Biotechnology Centre, Silesian University of Technology, Gliwice, Poland; Institute of Human Genetics, Polish Academy of Sciences, Poznan, Poland; University of Oxford, Department of Medicine, Medical Research Council Weatherall Institute of Molecular Medicine Centre for Computational Biology, Oxford, UK; Dana-Farber/Harvard Cancer Centre, Boston, MA 02215, USA; UMass Chan Medical School, MA 01605, USA; MD Anderson Cancer Center, The University of Texas, Department of Genetics, TX 77030, USA

**Author notes:** Correspondence /.

## Abstract

Oncogenes can be activated in *cis* through multiple mechanisms including enhancer hijacking events and noncoding mutations that create enhancers or promoters *de novo*. These paradigms have helped parse somatic variation of noncoding cancer genomes, thereby providing a rationale to identify noncanonical mechanisms of gene activation. Here we describe a novel mechanism of oncogene activation whereby focal copy number loss of an intronic element within the *FTO* gene leads to aberrant expression of *IRX3*, an oncogene in T cell acute lymphoblastic leukemia (T-ALL). Loss of this CTCF bound element downstream to *IRX3* (+224 kb) leads to enhancer hijack of an upstream developmentally active super-enhancer of the *CRNDE* long noncoding RNA (-644 kb). Unexpectedly, the *CRNDE* super-enhancer interacts with the *IRX3* promoter with no transcriptional output until it is untethered from the *FTO* intronic site. We propose that ‘promoter tethering’ of oncogenes to inert regions of the genome is a previously unappreciated biological mechanism preventing tumorigenesis.

## Introduction

The noncoding genome harbors differing classes of *cis*-reglatory elements including distal enhancers, poised promoters, and insulators that ensure precise control of gene expression across specialised tissues (1, 2). In cancer somatically acquired mutations of the noncoding genome can have deleterious *cis*-regulatory consequences by transcriptionally activating oncogenes through indels that generate *de novo* enhancers and promoters (3–10), by focal amplification of long-range enhancers (11, 12), through deletion of boundary elements (13–16), and structural rearrangements that lead to enhancer hijack (17–19). The pan-cancer analysis of whole genomes demonstrates that somatic mutation of the noncoding genome is a pervasive feature of cancer, yet variant interpretation is challenged through sequencing and variant calling limitations, and by the still nascent annotation of functional noncoding elements (20, 21). Addressing this requires the discovery of novel modes of oncogene activation, to simulatenously provide agency to noncoding mutations that may be overlooked as being *cis*-acting, and deliver broader insights into aberrant transcriptional regulation (22).

Previously, first-in-class mechanisms of oncogene activation following somatic mutation of the noncoding genome were discovered in T cell acute lymphoblastic leukemia (T-ALL), including *cis*-acting mutations that create a neomorphic enhancers, and recurrent focal deletions that disrupt boundaries between insulated neighbourhoods to activate *TAL1* and *LMO2* oncogenes respectively (6, 13). Analagous mechanisms were found to upregulate *FGD4* in bladder cancer, *ETV1* in colorectal cancer and *IRS4* in lung cancer (14, 23, 24). T-ALL exhibits archetypal cancer phenotypes including arrested cellular differentiation and replicative immortality of T cell progenitors following acquisition of somatic mutations (25–27). Additionally patients with refractory disease or those that relapse after induction chemotherapy have poor prognosis (28, 29). Although most T-ALL oncogenes *TAL1, LMO2, TLX1, TLX3, NKX2-1* and *MYB* are ectopically expressed by well characterized mechanisms of oncogene activation, some oncogenes are aberrantly expressed with no known genetic lesion, and thus provide an opportunity for the discovery of novel mechanisms of oncogene activation (30, 31).

In this study, we compared bulk RNA-seq between normal T cell subsets and T-ALL to identify *IRX3*, a putative oncogene in T-ALL with no known genetic driver (32). We show by ddPCR that a CTCF site residing within *FTO* intron 8 is recurrently deleted in T-ALL patients with concomitant expression of *IRX3*. We mimicked the deletion with CRISPR/Cas9 genome editing in a relevant *IRX3* negative T-ALL cell line to show that *FTO* intron 8 deletion is a causative step in the aberrant expression of *IRX3*. Next we utilised UMI4C to map chromatin looping about the *IRX3* locus, and discovered that *IRX3* is ‘tethered’ to the *FTO* intron 8 (+224 kb) whilst simultanouesly interacting with a developmental super-enhancer (-644 kb) overlapping *CRNDE*, but with no transcriptional consequence. We show that release of this ‘tether’ by focal deletion allows *IRX3* to hijack the *CRNDE* super-enhancer resulting in active transcription. We posit that ‘promoter tethering’ to inert regions of the genome is a previously unappreciated tumor suppressor mechanism, ensuring proto-oncogenes remain protected from activation by distal developmental super-enhancers and may explain the functional consequence of recurrent focal deletions in noncoding cancer genomes that as yet remain unexplained.

## Results

### *IRX3* is aberrantly expressed in T-ALL with enhancer-promoter contacts to neighbouring genes *FTO, CRNDE* and *IRX5*

We hypothesized that genes aberrantly expressed in T-ALL compared to their developmentally-matched normal cellular counterpart might uncover previously unrecognized oncogenes, enabling us to explore novel mechanisms of oncogene activation. To investigate this we analyzed gene expression obtained from normal hematopoietic stem cells (HSCs), common lymphoid precursors and six human thymic subsets isolated by FACS and subjected to RNA-seq (33). We generated a list of genes that were not expressed in any normal T cell subset (FPKM <0.1; n=2,468) and ranked their mean expression in 264 childhood cases of T-ALL (31). Within the top 50 aberrantly expressed gene list were well-characterized T-ALL oncogenes such as *TLX3, TLX1* and *NKX3-2*, validating this explorative approach (Fig. 1a). Our attention was drawn to *IRX3* (rank 20), given recent reports licencing it as an oncogene through its ability to immortalise HSCs in culture and induce T-lymphoid leukemias in vivo (32). *IRX3* encodes for an Iroquois-family homeobox transcription factor essential for normal organ development including limb bud pattern formation, nephron segmentation, and cardiac function, but with no known role in normal hematopoiesis (34–37).

**Fig. 1.**
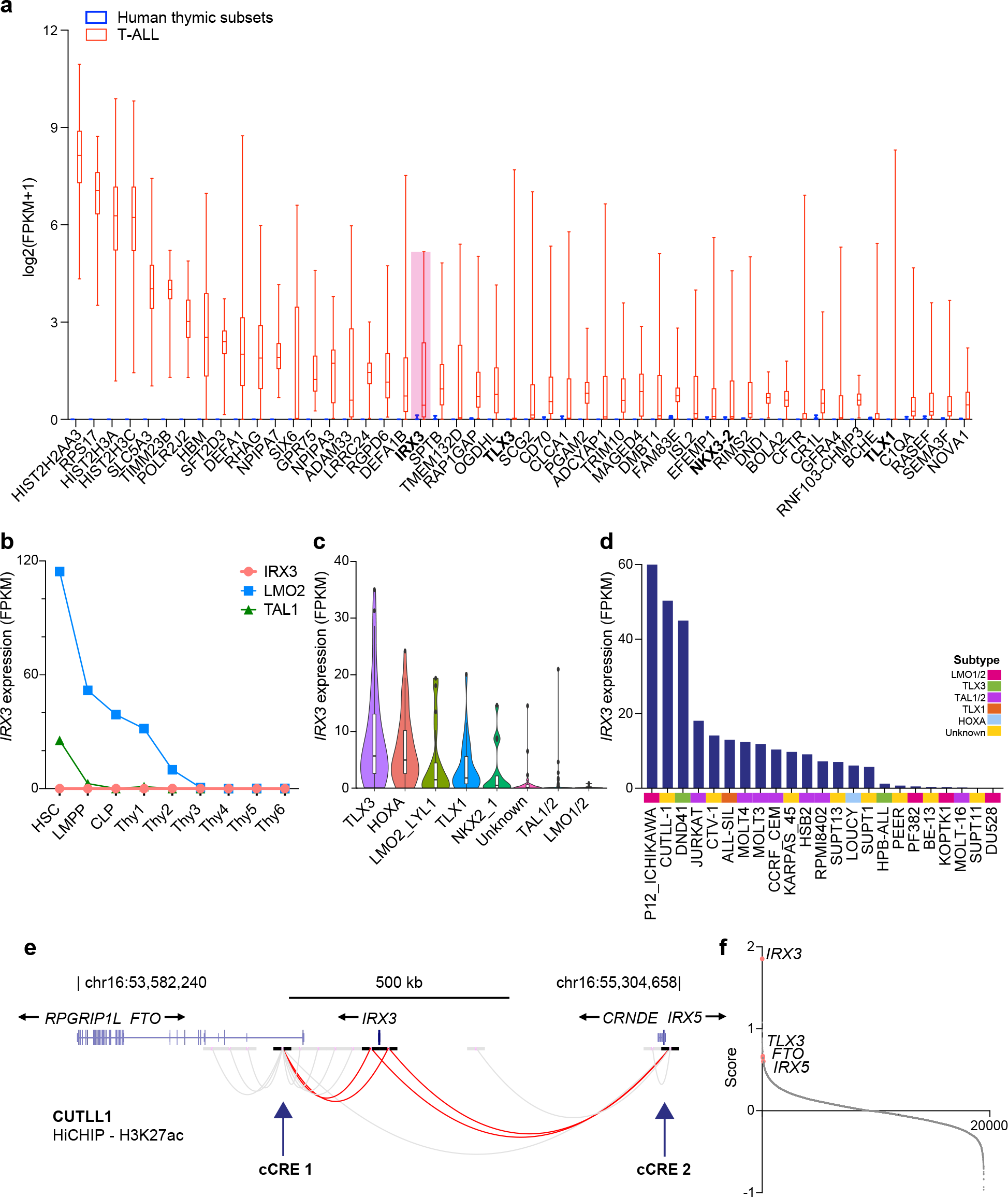
*IRX3* is aberrantly expressed in T-ALL and resides in a shared topologically associated domain with enhancer-promoter contacts to neighbouring genes *FTO, CRNDE* and *IRX5*. **a**, Box and whisker plot showing expression of top 50 genes aberrantly expressed in the St Jude’s paediatric T-ALL cohort (n=264) when compared to normal hematopoietic progenitors (NCBI GEO accession GSE69239); haematopoetic stem cells (HSCs), lymphoid primed multipotent progenitors (LMPPs), common lymphoid progenitors (CLPs) and T cell subsets (Thy1-6). Genes are ranked along the x-axis by median expression. **b**, Line graph tracking *IRX3, LMO2*, and *TAL1* expression by RNA-seq across hematopoietic and thymic progenitors. RNA-Seq from NCBI GEO accession GSE69239, with the following immunophenotypic definitions: from BM CD34^+^ cells, CD34^+^CD38^-^Lin^-^(HSCs), CD34^+^CD45RA^+^CD38^+^CD10^-^CD62L^hi^Lin^-^(LMPPs), CD34^+^CD38^+^CD10^+^CD45RA^+^Lin^-^(CLPs); from thymic CD34^+^ cells, CD34^+^CD7^-^CD1a^-^CD4^-^CD8^-^(Thy1), CD34^+^CD7^+^CD1a^-^CD4^-^CD8^-^(Thy2), and CD34^+^CD7^+^CD1a^+^CD4^-^CD8^-^(Thy3); from thymic CD34^-^cells, CD4^+^CD8^+^(Thy4), CD3^+^CD4^+^CD8^-^(Thy5), and CD3^+^CD4^-^CD8^+^(Thy6). **c**, Violin plot showing *IRX3* expression (FPKM) by RNA-seq from the St. Jude primary T-ALL cohort separated by class defining oncogenic subtypes (n=264). **d**, Bar chart showing *IRX3* expression (FPKM) by RNA-seq from T-ALL cell lines (n=24) and labelled by oncogenic subtype. **e**, Enhancer-promoter interactions about the *IRX3* locus mapped by HiChIP following pull-down for H3K27ac from the *IRX3* positive CUTTL1 T-ALL cell line (NCBI GEO accession GSE115896). Candidate cis-regulatory elements (cCREs) for *IRX3* are indicated with arrows. **f**, Ranked gene list by comparing *IRX3* positive (n=59) versus *IRX3* negative (n=59) T-ALL samples by RNA-seq from the St. Jude cohort. The y-axis ranking score metric for each gene was calculated by the GSEA ‘Signal2Noise’ computational method for categorical phenotypes (38, 39). Genes are listed along the x-axis in order of the ranked score.

In contrast to the developmental oncogenes *TAL1* and *LMO2, IRX3* is not expressed in any normal T-cell precursor (Fig. 1b). Further analysis of the pediatric T-ALL cohort separated according to their class-defining oncogenic subtypes showed that a greater proportion of the patients in the *TLX3* and *HOXA* subgroups had aberrant *IRX3* expression (defined as FPKM >1) when compared to the other subgroups (86% vs 23%; Fisher exact test statistic <0.00001, P <0.5) (Fig. 1c) (31). Furthermore, 74% (17/23) of T-ALL cell lines exhibited *IRX3* expression covering a wide spectrum of T-ALL subtypes, with P12-Ichikawa (*LMO2* subtype), DND-41 (*TLX3* subtype), and Jurkat (*TAL1* subtype) cells all displaying high levels of *IRX3* gene expression (Fig. 1d).

To date, no genetic drivers of aberrant *IRX3* expression have been identified in T-ALL. As the deregulated expression of genes can arise through *cis*-acting regulatory elements and/or disruption of topologically associated domains (TADs), we sought to map these structures about the *IRX3* locus. Analysis of published in-situ HiC data from normal human thymic tissue identified *IRX3* within a single TAD shared with *FTO, IRX5* and *CRNDE* encompassing 1.3 Mb (Supplementary Fig. 1, 2) (40, 41). To identify enhancer-promoter loops from *IRX3*, we next examined HiChIP data from the IRX3 positive CUTLL1 T-ALL cell line (Fig. 1e). This identified contacts 3’ to *IRX3* within *FTO* intron 8, and 5’ to *IRX3* within the *CRNDE* /*IRX5* locus. We named these two candidate *cis*-regulatory elements cCRE-1 and cCRE-2 respectively (Fig. 1e – red curved lines). These contacts suggested the transcriptional regulation of *IRX3* may originate intra-TAD through an undiscovered mechanism. Additionally, generation of a rank ordered gene list by comparing *IRX3* negative and *IRX3* positive primary T-ALL samples (n=118) by RNA-seq, identified the expression of multiple genes positively correlated with *IRX3* mRNA levels including *TLX3* at rank 57, *FTO* at rank 64, and *IRX5* at rank 109 out of 19,464 total genes (Fig. 1f). This provided further correlative evidence that cCRE-1 (*FTO*) and cCRE-2 (*CRNDE* /*IRX5*) may regulate *IRX3* expression through co-ordinated long-range interactions.

### *FTO* intron 8 is recurrently deleted in patients with T-ALL

We next explored whether copy number aberrations (CNAs) affected the *IRX3* CREs by examining copy number calls from published datasets of primary T-ALL patient samples and T-ALL cell lines. This analysis revealed 13 T-ALL genomes (12 of patient origin and 1 cell line – ALL-SIL) with heterozygous copy number losses impinging on the *FTO* gene, and notably all intersected with cCRE-1 (Fig. 2a). No significant CNAs were identified at the cCRE-2 – *CRNDE* /*IRX5* locus or the *IRX3* locus itself. Expression data was available for 5/13 T-ALL samples and these exhibited aberrant expression of *IRX3* mRNA, with 1 primary patient sample having allele specific expression (ASE) indicative of a heterozygous *cis*-acting genetic lesion (Fig. 2b). Due to relatively small size of *IRX3* exon sequences (2.6 kb total length), the gene often lacks informative heterozygous SNPs to make consistent ASE calls across T-ALL genomes. To overcome this challenge, we used the CCLE promoter methylation dataset covering 15 T-ALL cell lines, to identify promoter methylation allelic imbalance (0.4<promoter-methylation^IRX3^<0.6) as a proxy for allele-specific gene expression (Supplementary Fig. 3). This approach effectively identified ALL-SIL as cell line with potential ASE, and indeed this cell line not only had the aforementioned copy number loss, but also associated supraphysiological expression of *IRX3* (Fig. 2a, b).

**Fig. 2.**
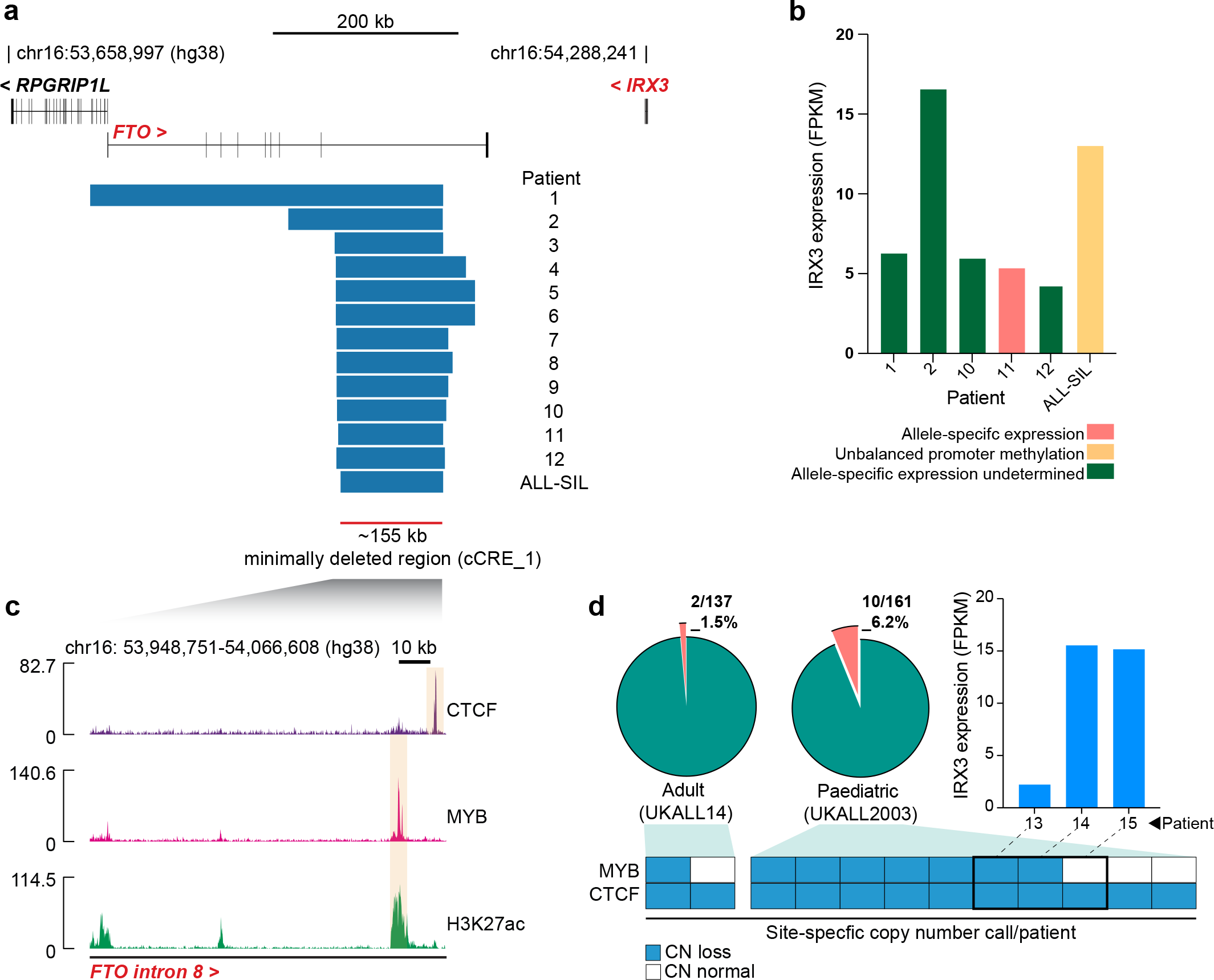
Recurrent deletions of *FTO* intron 8 in T-ALL patients impinge on a CTCF binding site. **a**, Focal heterozygous deletions identified in T-ALL genomes at *FTO*. 12 originate from primary patient samples and 1 from the ALL-SIL T-ALL cell line. Deletions identified in patient samples are from multiple cohorts including St. Jude, UKALL2003, and ICGZ Poznan **b**, *IRX3* expression (FPKM) by RNA-seq from matched primary patient samples and the ALL-SIL T-ALL cell line harbouring *FTO* intron 8 deletions. Allele-specific expression identified in patient 11 was determined by St. Jude’s prior analyses, and unbalanced promoter methylation for the ALL-SIL cell line was ascertained by analysis of the CCLE Promoter Methylation dataset. **c**, ChIP-Seq for CTCF (NCBI GEO accession GSE68976), MYB and H3K27ac (NCBI GEO accession GSE76783) in the Jurkat T-ALL cell line centred on the minimally deleted region within *FTO* intron 8. **d**, Pie charts showing the frequency of *FTO* intron 8 deletions determined by ddPCR in adult (n=137) and paediatric (n=161) T-ALL cohorts. Stacked boxes summarise CTCF or MYB site-specific copy number calls for each patient identified with *FTO* intron 8 deletions and a bar chart showing *IRX3* expression (FPKM) is shown for patient samples where matched RNA was available for sequencing.

Interestingly, the copy number losses mapped to a minimally deleted region ( 155 kb) within *FTO* intron 8, alluding to a sequence element within this intron with regulatory capacity (Fig. 2a). To fine-map sequences with regulatory capacity, we examined previously generated ChIP-Seq datasets from the *FTO* intron 8 wild-type Jurkat T-ALL cell line. We rationally selected specific ChIP-Seq datasets to explore, including the oncogenic helix-turn-helix transcription factor MYB, the enhancer histone mark H3K27ac, and the TAD-boundary/chromatin looping transcription factor CTCF, as these have been previously implicated in aberrant gene expression mechanisms of T-ALL (6–9, 31, 42). The minimally deleted region was largely devoid of high-amplitude ChIP-Seq peaks, except for a single CTCF binding peak, and a single MYB peak enriched with H3K27ac. From this we hypothesised that loss of CTCF and/or MYB binding within *FTO* intron 8 may be involved in the deregulated expression of *IRX3* (Fig. 2c).

To explore this hypothesis further, we developed a digital droplet PCR (ddPCR) assay with hydrolysis probes capable of picking up copy number alterations at the CTCF or MYB binding sites within *FTO* intron 8 (Supplementary Fig. 4). This had two aims, first to ascertain the frequency of *FTO* intron 8 copy number aberrations (CNAs) in a larger cohort of primary T-ALL samples, and secondly to allow for independent copy number calls at both loci where differences between the calls may provide mechanistic insight. Among 298 unselected primary T-ALL samples collected at diagnosis, CNAs within *FTO* intron 8 were identified in 2/137 (1.4%) adult and 10/161 (6.2%) pediatric samples of T-ALL, demonstrating enrichment for *FTO* intron 8 CNAs in the pediatric setting (p = 0.04 by Chi-Squared test, Fig. 2d).

While 8/12 had copy number loss for both the CTCF and MYB sites, notably 4/12 had heterzygous copy number loss of the CTCF site alone, suggesting that loss of this CTCF site is most likely to be functionally relevant. Furthemore, we identified aberrant *IRX3* expression in 3/12 samples with *FTO* intron 8 CNAs by analysis of patient matched RNA-seq (Fig. 2d). RNA-seq data was unavailable for the remaining (9/12) samples.

### CRISPR/Cas9 mediated disruption of *FTO* intron 8 CTCF site transcriptionally activates *IRX3*

Following identification of recurrent copy number losses in primary T-ALL samples, we sought to determine if deletion of *FTO* intron 8 CTCF and MYB binding sites, either individually, or together, is functionally capable of upregulating expression of *IRX3*. To address this, CRISPR/Cas9 was used to introduce indels at the CTCF and MYB binding sites in the PF-382 T-ALL cell line (Fig. 3a). This cell line is an ideal model for this experiment as it is negative for *IRX3* mRNA (FPKM = 0.49, Fig. 1d), it harbors comparable enhancer architecture across the *FTO*/*IRX3*/*CRNDE* /*IRX5* genomic loci as other *IRX3* positive T-ALL cell lines, and has normal copy number at *FTO* intron 8 (Supplementary Fig. 5, Supplementary Table. 1). Following single cell sorting and expansion, colonies were genotyped by flanking PCR (Supplementary Fig. 6). Deletions of 12kb that impinged on both CTCF and MYB binding sites, thus mimicking the copy number losses observed in the primary patient samples, led to upregulation of *IRX3* mRNA relative to the parental PF-382 cell line and unedited single cell derived clones (Fig. 3b).

**Fig. 3.**
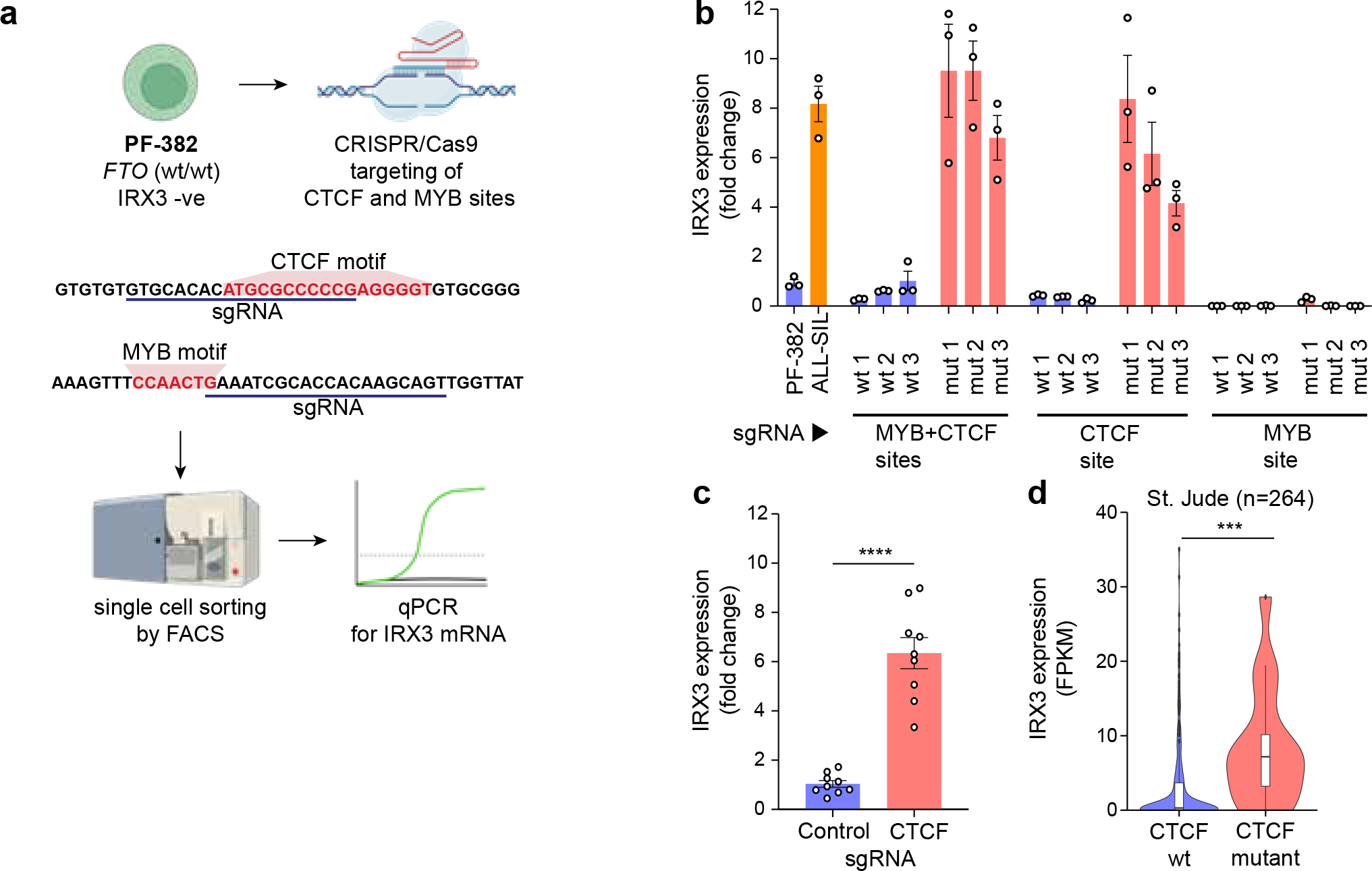
Deletion of *FTO* intron 8 CTCF site causes transcriptional activation of *IRX3*. **a**, Experimental outline for CRISPR/Cas9-mediated disruption of *FTO* intron 8 CTCF and MYB sites in the PF-382 (FTO wt/wt) and *IRX3* negative (FPKM<1) T-ALL cell line. **b**, Bar chart showing *IRX3* expression determined by qPCR for PF-382 (*FTO* wild-type) and ALL-SIL (*FTO* intron 8 deleted) T-ALL cell lines and unedited (wt) and edited (mut) clones. Data presented are 3 technical replicates +/- SD for each biological sample. **c**, Bar chart showing *IRX3* expression determined by qPCR for PF-382 polyclonal edited cells following CRISPR/Cas9 mediated disruption of the *FTO* intron 8 CTCF site. Technical replicates from 3 independent experiments are shown. **d**, Bar chart showing *IRX3* expression (FPKM) of primary T-ALL samples with (n=16) and without (n=248) CTCF mutations from the St. Jude T-ALL cohort. p=0.0007 from two-tailed T test.

Similar upregulation of *IRX3* mRNA was observed in clones with sole disruptive indels of the CTCF binding site, but crucially not observed following indels that affect the MYB site alone. Furthermore, deletion of the CTCF binding site in a polyclonal population led to a significant increase of *IRX3* mRNA relative to unedited controls (Fig. 3c). These data, in addition to the prior observation of copy number losses in patient samples only affecting the CTCF site, suggests that loss of CTCF binding within *FTO* intron 8 is causal for *IRX3* mRNA upregulation in a subset of T-ALL. As *FTO* intron 8 copy number aberrations are not pervasive across all *IRX3* positive T-ALL samples, we postulated that mutations of CTCF itself may create the same phenotypic outcome of elevated *IRX3* expression. To explore this, we compared RNA-seq data from CTCF wild-type and CTCF mutant T-ALL samples, and found significantly higher *IRX3* expression in the CTCF mutant group (Fig. 3d, Supplementary Table 2). This correlatively suggests that mutations that affect CTCF function may have a consequence on *IRX3* gene regulation, and may provide an explanation for aberrant *IRX3* expression in lieu of the aforementioned *FTO* intron 8 copy number aberrations, particularly in TLX3 positive T-ALL cases where CTCF mutations are most frequent.

### Focal deletion of the *IRX3* ‘promoter tether’ residing within *FTO* intron 8 allows enhancer hijack

Given the importance of CTCF as an architectural protein with fundamental roles in chromatin looping, and its ability to cluster at TAD boundaries, we hypothesised that loss of CTCF binding within *FTO* intron 8 may allow previously non-permissive interactions between the *IRX3* promoter and distal enhancer elements. To investigate this hypothesis, we quantified interactions from the *FTO* intron 8 CTCF site and *IRX3* promoter by UMI-4C. Baiting the *FTO* intron 8 CTCF site in PF-382 (IRX3 negative) cells revealed a dense cluster of looping interactions between the CTCF site and *IRX3*, suggesting that the proximal promoter of *IRX3* is tethered to the *FTO* intron 8 in the wild-type setting (Fig. 4a). By comparing ALL-SIL cells (IRX3 positive) harboring the *FTO* intron 8 deletion and PF-382 cells (IRX3 negative) wild-type for *FTO*, we discovered a marked increase in the number of interactions between the *IRX3* promoter bait and the *CRNDE* locus in ALL-SIL cells, previously identified as cCRE-2 (Fig. 4a, b).

**Fig. 4.**
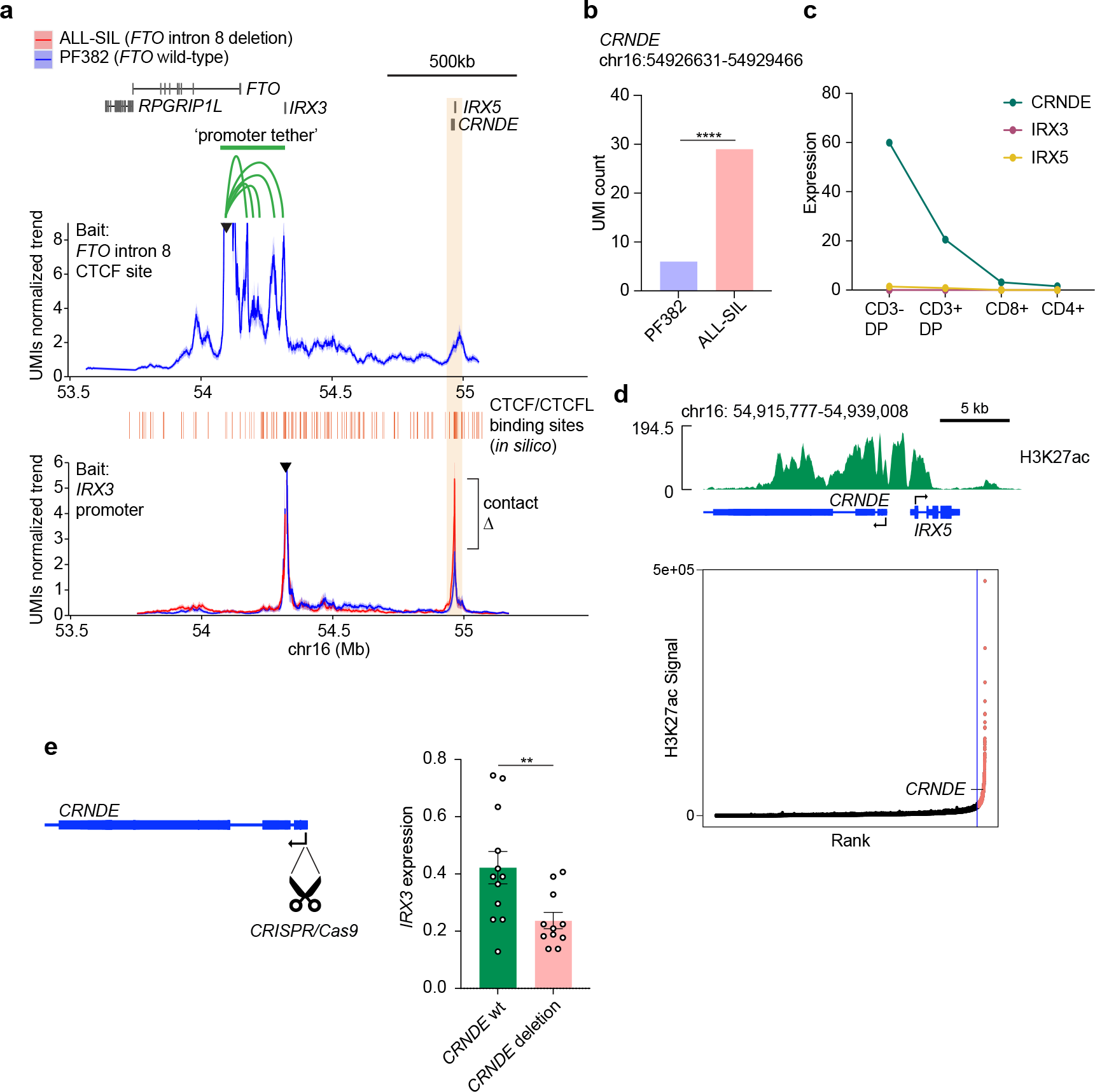
Transcriptional activation of *IRX3* in T-ALL by loss of a promoter tether and enhancer hijack from the *CRNDE* locus. **a**, (Top panel) UMI-4C contact profile generated by baiting the *FTO* intron 8 CTCF site in PF382 (*FTO* wild type) cells. A green bar highlights the contacts that form a promoter tether between IRX3 and *FTO* intron 8. (Bottom panel) UMI-4C contact profile generated by baiting the *IRX3* proximal promoter in the PF382 (*FTO* wild type) and ALL-SIL (*FTO* intron 8 heterozygous deleted) T-ALL cell line. **b**, UMI count of interactions between the *IRX3* proximal promoter and the *CRNDE* locus (p = 0.0005; Fisher’s exact test; UMI4Cats analysis package). **c**, Expression (FPKM) of *CRNDE, IRX5* and *IRX3* across T cell subsets by RNA-seq. **d**, PF-382 cell line H3K27ac ChIP-Seq profile (NCBI GEO accession GSE76783) at the *CRNDE* /*IRX5* locus (top) and rank ordering of super-enhancers (ROSE) analysis (red dots indicate regions classed as super-enhancers). **e**, Expression level of *IRX3* mRNA as determined by qPCR from ALL-SIL T-ALL cell line following CRISPR/Cas9 mediated deletion of the *CRNDE* super-enhancer locus.

Interestingly, high levels of *CRNDE* expression (FPKM>20) are observed in developing CD3^-^ double positive (DP) and CD3^+^ DP T cells, with lower but maintained expression in mature single positive CD8^+^ and CD4^+^ T cells, meaning this locus is actively transcribed during normal T cell development (Fig. 4c). Further examination of the *CRNDE* locus by H3K27ac ChIP-Seq in PF-382 cells identified an enhancer region covering the proximal promoters of *CRNDE* and *IRX5*, in addition to the entire length of the *CRNDE* gene body (Fig. 4d). Notably, rank ordering of all PF-382 enhancers by the ROSE algorithm further classified this locus as a super-enhancer (113th out of 23,737 total enhancers; Fig. 4d).

Indeed, disruption of the *CRNDE* enhancer in ALL-SIL cells using CRISPR/Cas9 resulted in two-fold downregulation of *IRX3* expression (p-value = 0.01; Fig 4e). These observations confirm that aberrant *IRX3* expression following deletion of the *FTO* intron 8 CTCF binding site is driven by increased interaction with the *CRNDE* /*IRX5* super-enhancer.

## Discussion

In this study we discovered recurrent focal deletions of a *cis*-regulatory element within *FTO* intron 8 for the oncogene *IRX3* in T-ALL. Although situated downstream to *IRX3*, deletion of this long-range insulator counterintuitively permits enhancer hijack of an upstream super-enhancer. This is distinct from canonical TAD fusion events in cancer whereby focal deletions or methylation disrupts boundary elements positioned between the oncogene and *cis*-regulatory effector (13, 14, 16, 43). In contrast to previously discovered enhancer hijack events, where the enhancer-promoter (E-P) interaction remains naïve until the structural rearrangement occurs, we reveal interactions between the *IRX3* promoter and *CRNDE* super-enhancer in *IRX3* negative cells (17–19). We posit that the *IRX3* promoter is sequestered to a relatively inert region of *FTO* intron 8, yielding minimal transcriptional output, despite residual interactions with the *CRNDE* super-enhancer (Fig. 5). This sequestration is facilitated by CTCF binding at the *FTO* intron leading to the formation of a ‘promoter tether’. Therefore, loss of this CTCF site by focal deletion untethers the *IRX3* promoter, providing an example of enhancer-promoter competition occurring within the same TAD (Fig. 5). We thus demonstrate a new somatically acquired mechanism of *IRX3* activation initiated through somatic deletions of *FTO* intron 8, distinct from recent findings in acute myeloid leukemia (AML) describing an intronic long noncoding RNA in *FTO* that regulates *IRX3* expression (44). Together, these findings add to the complex regulatory relationship between the *FTO* and *IRX3* genes first identified through the discovery of obesity-associated germline variants (45, 46). We further speculate that promoter tethering to inert regions of the genome is a previously unappreciated tumor suppressor mechanism through which potent oncogenes are protected from activation of nearby developmental super-enhancers. Integrating 3-dimensional promoter interactions with copy number data may highlight further examples of this phenomenon, potentially explaining the functional consequence of recurrent focal deletions in noncoding genomes of cancers that as yet remain unexplained.

**Fig. 5.**
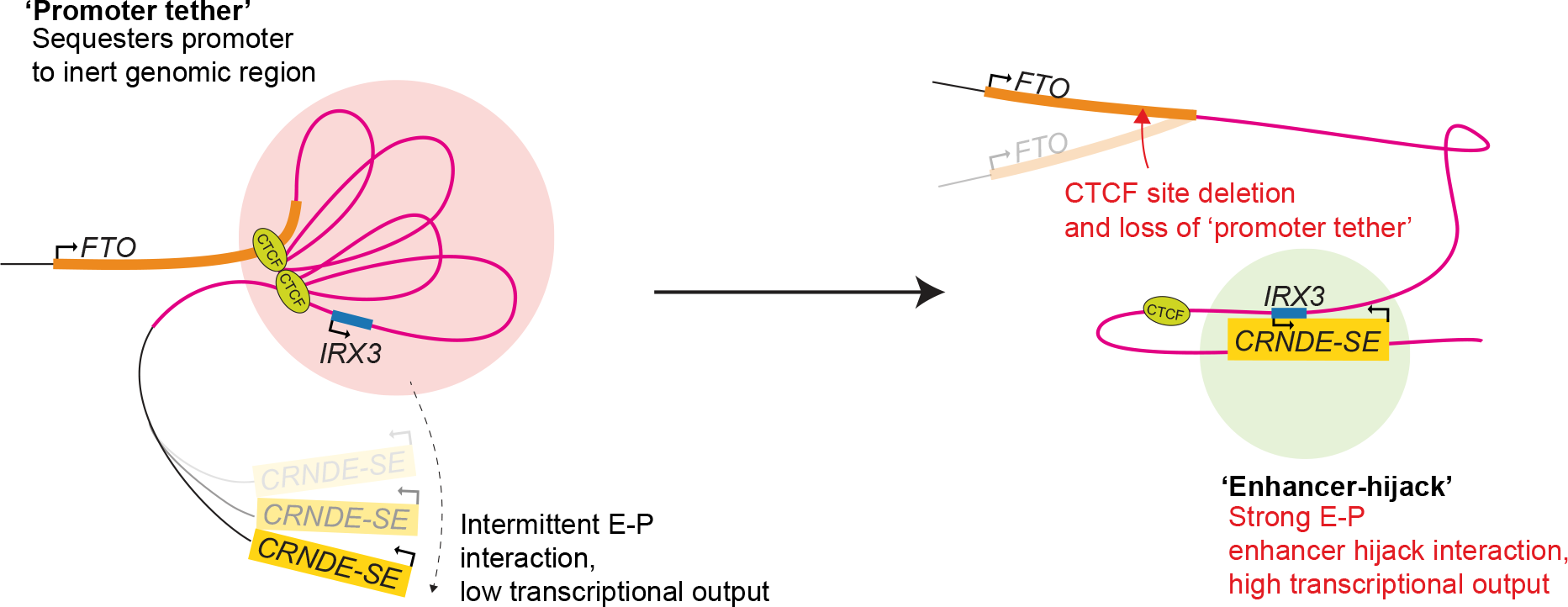
Mechanism of *IRX3* transcriptional activation in T-ALL. Proposed mechanism of action whereby promoter tethering of *IRX3* to the relatively inert *FTO* intron 8 locus by CTCF binding allows infrequent interaction enhancer-promter interactions and low transcriptional output. Subsequent focal deletion of this intronic CTCF site leads to loss of the promoter tether and enhancer hijack of the *CRNDE* super-enhancer.

## Methods

### Digital droplet PCR (ddPCR)

Identification of focal copy number aberrations within *FTO* intron 8 was achieved by ddPCR. Probes were designed by using the Bio-Rad Assay Design Engine (https://www.bio-rad.com/digital-assays/assays-create/cnd) against two loci of interest and one reference locus with the following co-ordinates; CTCF binding site hg19|chr16:54096903-54097025 with probe fluorophore FAM, and MYB binding site hg19|chr16:54084994-54085116 with probe fluorophore HEX, and reference region *AP3B1* hg19|chr5:77358602-77358724 with probe fluorophore HEX. All ddPCR reactions were set up as per the following, 11μL 2x ddPCR Supermix (1863023), 0.36μL 20x MYB-HEX, 0.74μL 20x AP3B1-HEX, 1.1μL 20x CTCF-FAM, 1.1μL HaeIII (NEB – R0108S), 1 μL patient sample gDNA with a concentration range of 10 – 66 ng/μL and 6.7 μL nuclease free water to make a 20 μL total reaction. Droplets were generated on a QX200 Droplet Generator (1864003) as per manufacturers’ instructions and plates were sealed using a PX1 PCR Plate Sealer (1814000). Droplets were transferred to a thermocycler for PCR with the following parameters; 1 cycle of 95C for 10 mins, 40 cycles of 94C for 30 sec followed by 60C for 1 min, 1 cycle of 98C for 10 min. Droplets were then read on a QX200 Droplet Reader (1864003) and copy number calls for the two loci of interest, for each sample, was determined by using the QuantaSoft Analysis Pro (Bio-Rad) software.

### CRISPR/Cas9 genome editing of PF-382 cell line

Deletion and/or mutation of MYB and CTCF binding sites of *FTO* intron 8, singly and together was achieved by CRISPR/Cas9 genome editing. A tripartite ribonucleoprotein (RNP) delivery system composed of recombinant Cas9 (IDT - 1081058), tracrRNA-ATTO550 (IDT - 1075927) and custom designed crRNA (IDT) targeting the genomic loci of interest was used. To target the MYB binding site the crRNA sequence (guide RNA) used was 5’-GAAATCGCACCACAAGCAGT-3’, to target the CTCF binding site the crRNA sequence used was 5’-GTGCACACATGCGCCCCCGA-3’, and to target both sites simultaneously both aforementioned crRNAs were mixed at a 1:1 ratio for complex formation with the tracrRNA-ATTO550. Ribonucleoprotein complexes were electroporated into PF-382 cells at a cellular density of 2M cells/electroporation with 10 μL of Cas9:crRNA:tracrRNA-ATTO550 complex, 3 μL IDT electroporation enhancer (IDT - 1075915), and 100 μL Ingenio Electroporation Solution (Mirus - MIR50111). Samples were electroporated using the D-23 program on an Amaxa Nucleofector. Cells were allowed to recover for 48 hours prior to single cell sorting by flow cytometry into 96 well plates. Cells were sorted by ATTO550 positivity, where the top 5% of ATTO550 positive cells were sorted and incubated under standard tissue culture conditions (37C and 5% CO2) in RPMI supplemented with 10% FCS and 5% pen/strep. The plates were examined for colonies 3 weeks after sorting, and gDNA was extracted by using QuickExtract DNA Extraction solution (Epicentre) as per manufacturer’s instructions. Clones were screened for mutations by PCR with primers flanking the target editing sites. Primers used to identify indels following editing of the MYB binding site were 5’- TGCTCCAACAAATCAACAGAACT-3’ (forward) and 5’- TGGAACTCAAGGCACAGAAAAC-3’ (reverse). Primers used to identify indels following editing of the CTCF binding site were 5’-GTGAAAGTGGCTGGGGATCA-3’ (forward) and 5’-AGACACAACATCCACAGTTCT-3’ (reverse). For PCRs we used Phusion High-Fiedlity PCR Master Mix with HF Buffer (NEB) and reactions were run as per manufacturer’s instructions. Primers used to identify deletion of both MYB and CTCF binding sites together were 5’- ATGGATGAAGAGGAAGAGAAGGTTC-3’ (forward) and 5’-CTTCTAAACCAAATGTGCCTGAACT-3’ (reverse). To ensure simultaneous amplification of both the large unedited ( 12.6 kb) and edited allele (646 bp) we used the LongAmp Taq DNA Polymerase (NEB) which is specially formulated to amplify large products. Reactions were set up as per manufacturer’s instructions. Deletion of the *FTO* intron 8 CTCF binding site in PF-382 cells to generate a polyclonal edited population was achieved as per the above, by using the crRNA sequence targeting the CTCF binding site 5’-GTGCACACATGCGCCCCCGA-3’ followed by bulk sorting by FACS of the ATTO550 positive population.

### CRISPR/Cas9 genome editing of ALL-SIL cell line

Deletion of the *CRNDE* super-enhancer was achieved by CRISPR/Cas9 genome editing using the aforementioned tripartite ribonucleoprotein (RNP) delivery system composed of recombinant Cas9 (IDT - 1081058), tracrRNA-ATTO550 (IDT - 1075927) and custom designed crRNAs (IDT). The cRNA sequences used were as follows 5’-TCTCGATCGCGCTATTGTCA-3’ and 5’-AGCCCACGGGACGTCTGGTC-3’ and were designed to target the transcriptional start site of the *CRNDE* noncoding RNA to collapse the *CRNDE* super-enhancer. RNA duplex formation was achieved by combining 0.75 μL of each crRNA at a 1:1 ratio or 1.5 μL of negative crRNA with 1.5 μL tracrRNA-ATTO550, followed by heating in a thermocycler to 95C for 5 mins and cooling to room temperature. The cas9:crRNA:tracrRNA-ATTO550 complex was formed by adding 3 μL of HiFi Cas9 followed by a 10 min incubation at room temp. Ribonucleoprotein complexes were electroporated with the Neon Transfection System (Thermo Fisher Scientific) as per manufacturer’s instructions at a cellular density of 0.6 x106 cells/electroporation with 10μL Neon Transfection tips. Electroporation conditions used were 1700V/20ms/1 pulse. After 48 hours, cells were bulk sorted by FACS by gating on the ATTO550 positive fraction. Deletion of the *CRNDE* super-enhancer was validated by PCR by using the following primers 5’-GTGACAGACTCGGGTTCTCG-3’ (forward) and 5’- TCACCTATGATTGGGCGCTG-3’ (reverse). PCRs reactions used KAPA HiFi HotStart ReadyMix (Roche) as per manufacturer’s instructions using an annealing temperature of 65.7C.

### Quantitative PCR (qPCR) of *IRX3*

Total RNA was extracted from samples by using a RNease Mini Kit (Qiagen) by following manufacturer’s instructions including on column DNase digestion, and concentrations were measured on a Nanodrop One (ThermoFisher). A total of 1 μg of RNA was used as input for cDNA synthesis reactions by using the Omniscript RT Kit (Qiagen) along with random hexamers. All qPCR reactions used FastStart Universal Probe Master (Merck) with the manufacturer’s recommended reaction conditions with some minor adjustments. Reactions were run with 5% DMSO spike-in and cycling conditions were adjusted to the following; 1 cycle of 95ºC for 10 min, followed by 40 cycles of 95ºC for 15s and 60.8ºC for 1 min. Samples were run in a 96 well plate format on a Mastercycler epgradient S thermocycler (Eppendorf). Primer pairs to quanitify to IRX3 mRNA were 5’-CTCACAGACTGGTCTCAGCG-3’ (forward), 5’-TCTGCCCCTCCGCACCTGCT-3’ (FAM probe), and 5’-AGGCACTACAGCGATCTGTTC-3’ (reverse). The primers pairs for the housekeeping gene GAPDH were 5’-TGCACCACCAACTGCTTAGC-3’ (forward), 5’-ACCCCTGGCCAAGGTCATCCATGA-3’ (FAM probe) and 5’-GGCATGGACTGTGGTCATGAG-3’ (reverse) Normalised expression ratios were calculated by the efficiency corrected Ct method whilst using GAPDH as the endogenous reference mRNA as described at length by Bookout et al (47).

### UMI-4C

Interactions between the *FTO* intron 8 CTCF site and IRX3 proximal promoter with distal loci were identified by UMI-4C as described previously with the following minor adjustments (48). Briefly, 5 μg of 3C template from each experimental condition was sonicated at 4ºC for 20 cycles (30s ON/ 60 s OFF) on a Bioruptor Pico (Diagenode). The post-sonication size distribution was checked on a 2200 Tapestation (Agilent) with D1000 Screentape to ensure an ideal peak distribution size of 450 – 550 bp. The sonicated DNA was end-repaired by the End Repair Module (NEB – E6050L) and cleaned up with 2.2x AmpureXP beads (Beckman Coulter) followed by dA-tailing (NEB – E6053L) as per the manufacturers’ reaction conditions and instructions. Following A-tailing, DNA was cleaned up with 2x AmpureXP beads and eluted in 67 μL of EB buffer (Qiagen). Illumina compatible adapters (IDT – xGEN UDI-UMI adapters) were ligated by using T4 ligase (NEB – M0202S) overnight at 16ºC. Two nested PCR reactions were conducted with the following bait sequences for the *FTO* intron 8 CTCF site, 5’-GAAAGGCTTGTTCTTCCTGGC-3’ for bait 1 (upstream primer), 5’-AATGATACGGCGACCACCGAGATCTACACTCTTT CCCTACACGACGCTCTTCCGATCTTGGAAC TGCTGTTTCCCACTT-3’ for bait 2 (Illumina prefix appended to downstream primer). The bait sequences for the IRX3 proximal promoter were, 5’-AGGAAGAAGTGAAGGGAACGGA-3’ for bait 1 (upstream primer), 5’-AATGATACGGCGACCACCGAGATCTACACTCTTT CCCTACACGACGCTCTTCCGATCTTGCAGGA GCCCGAAGCA-3’ for bait 2 (Illumina prefix appended to downstream primer) and, 5’-CAAGCAGAAGACGGCATACGA-3’ for the Illumina enrichment primer. The PCR reactions were performed in 50 μl volume with the following reagents: 10 μl GoTaq Flexi buffer (Promga M792A), 3 μl MgCl 25 mM (Promega A3511), 0.2 mM dNTPs, 2 μl 10 mM bait 1/upstream primer (0.4 mM final concentration), 2 μl 10 mM Illumina enrichment primer (0.4 mM final concentration), 1 μl GoTaq Hot Start (Promega M5005) and 200 ng UMI-4C DNA template. PCR program: 2 min 95 °C, 20 cycles of 30 s 95 °C, 30 s 56 °C and 60 s 72 °C and final extension of 5 min 72 °C. Amplified DNA was cleaned with 1x AmpureXP beads, and the eluate was used as a template in the second PCR reaction. Conditions for the second PCR reaction were identical to the first, and the only differences were the use of the bait primer 2. The molarity of the libraries were quantified by using a Qubit dsDNA HS Assay Kit (ThermoFisher – Q32851) together with a Tapestation (Agilent). Libraries were pooled and diluted to 4 nM concentration and multiplexed with other libraries, including a 10% PhiX spike-in (Illumina) and sequenced on a MiSeq (Illumina) with a MiSeq Reagent Nano Kit v2, 300 cycles (Illumina). FASTQs generated were analysed with the ‘UMI4Cats’ R package to generate contact maps against the IRX3 proximal promoter (49).

### Primary samples

Diagnostic genomic DNA was available from 161 paediatric and 137 adult patients with previously untreated T-ALL who were enrolled onto the UKALL2003 and UKALL14 trials respectively. UKALL2003 is registered under the ISRCTN number 07355119 at http://www.controlled-trials.com and was initially opened in 2003 for patients aged 1-18 years (50). In 2006 the upper age limit of the trial was increased to 20 years, and in 2008 it was increased to 25 years of age. Ethical approval for the trial was obtained previously from the Scottish Multi-Centre Research Ethics Committee on 25/02/2003, ref: 02/10/052. UKALL14 is registered with ClincialTrials.gov, number NCT01085617, and has the ISRCTN number 66541317. Adult patients aged 25-65 years old with newly diagnosed T-ALL were enrolled onto this trial between 2012 and 2018. Ethical approval was previously obtained from the London-Fulham Research and Ethics Committee, ref: 09/H0711/90. All samples from both UKALL2003 and UKALL14 trials were collected from patients with informed consent according to the Declaration of Helsinki.

## ACKNOWLEDGEMENTS

The authors thank the patients, families, and clinical teams who have been involved in all trials. Primary childhood leukemia samples used in this study were provided by VIVO Biobank, supported by Cancer Research UK Blood Cancer UK (Grant no. CRCPSC-Dec2100003). Samples were acquired with support from laboratory teams in the Bristol Genetics Laboratory, Southmead Hospital, Bristol, United Kingdom; Molecular Biology Laboratory, Royal Hospital for Sick Children, Glasgow, United Kingdom; Molecular Haematology Laboratory, Royal London Hospital, London, United Kingdom; and Molecular Genetics Service and Sheffield Children’s Hospital, Sheffield, United Kingdom. This work was supported in part by Cancer Research UK CRUK/A13920 to A.K.F for UKALL14 trial and CRUK/A21019 to A.K.F for UKALL14 Biobank. S.R was funded by a John Goldman Fellowship from Leukaemia UK (2019-2022). M.R.M. is supported through a GOSH Children’s Charity Professorship. This work was supported by the Francis Crick Institute which receives its core funding from Cancer Research UK (CC2008), the UK Medical Research Council (CC2008), and the Wellcome Trust (CC2008). For the purpose of Open Access, the authors have applied a CC BY public copyright license to any Author Accepted Manuscript version arising from this submission. P.V.L. is a CPRIT Scholar in Cancer Research and acknowledges CPRIT grant support (RR210006). M.D and P.V.V were supported by the European Union’s Horizon 2020 research and innovation program under Grant agreement no. 952304. This project was supported by funding from The National Centre for Research and Development: STRATEGMED3/304586/5/NCBR/2017. S.R. would like to thank A. Motazedian, A. Das, E. Wainwright and D. Vassiliadis for academic discussions pertaining to this study and Lorna Neal for invaluable non-academic discussions.

