## Supplementary Figs 1-6, Supplementary Table 1-2 for "Focal Deletions of a Promoter Tether Activate the *IRX3* Oncogene in T Cell Acute Lymphoblastic Leukemia"

Rahman et al.

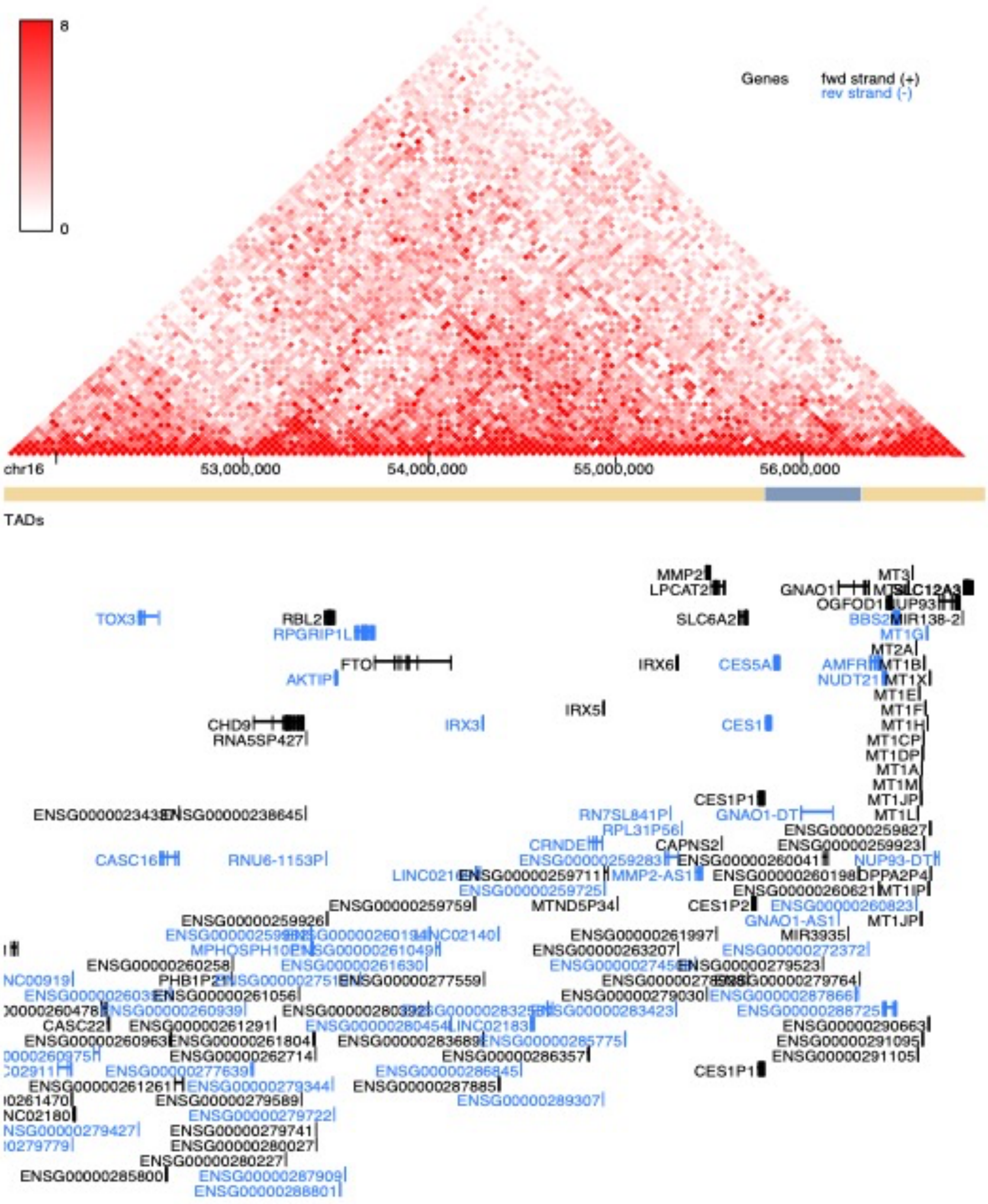


**Supplementary Figure 1. HiC data at 40 kb resolution from normal human thymic tissue.** Heatmap data about the *IRX3* locus generated from normal human thymic tissue. Experimental data was generated by Leung et al. 2015 with the NCBI GEO accession #GSE58752. Alternating bars coloured brown>blue>brown underneath the heatmap represent topological associated domains (TADs). Genomic co-ordinates are in hg38 and data was downloaded from http://3dgenome.fsm.northwestern.edu/ (last accessed on 24th August 2023).

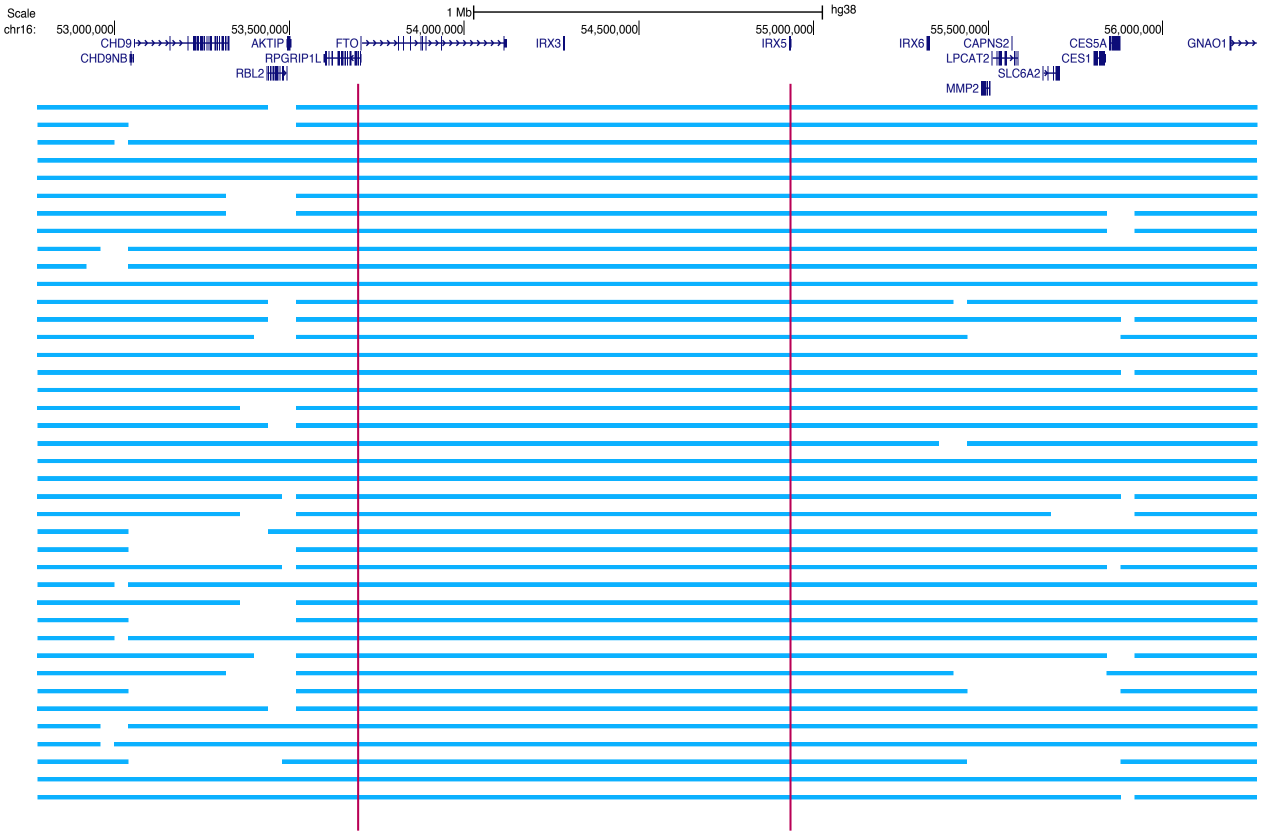


**Supplementary Figure 2. Topologically associated domains (TADs) from 40 human tissues about the *FTO*/*IRX3*/*IRX5* locus.**

TADs are shown as horizontal light blue lines and the *FTO*/*IRX3*/*IRX5* locus is demarcated by two vertical red lines. Genomic co-ordinates are in hg38 and data was downloaded from http://3dgenome.fsm.northwestern.edu/ (last accessed on 24th August 2023).

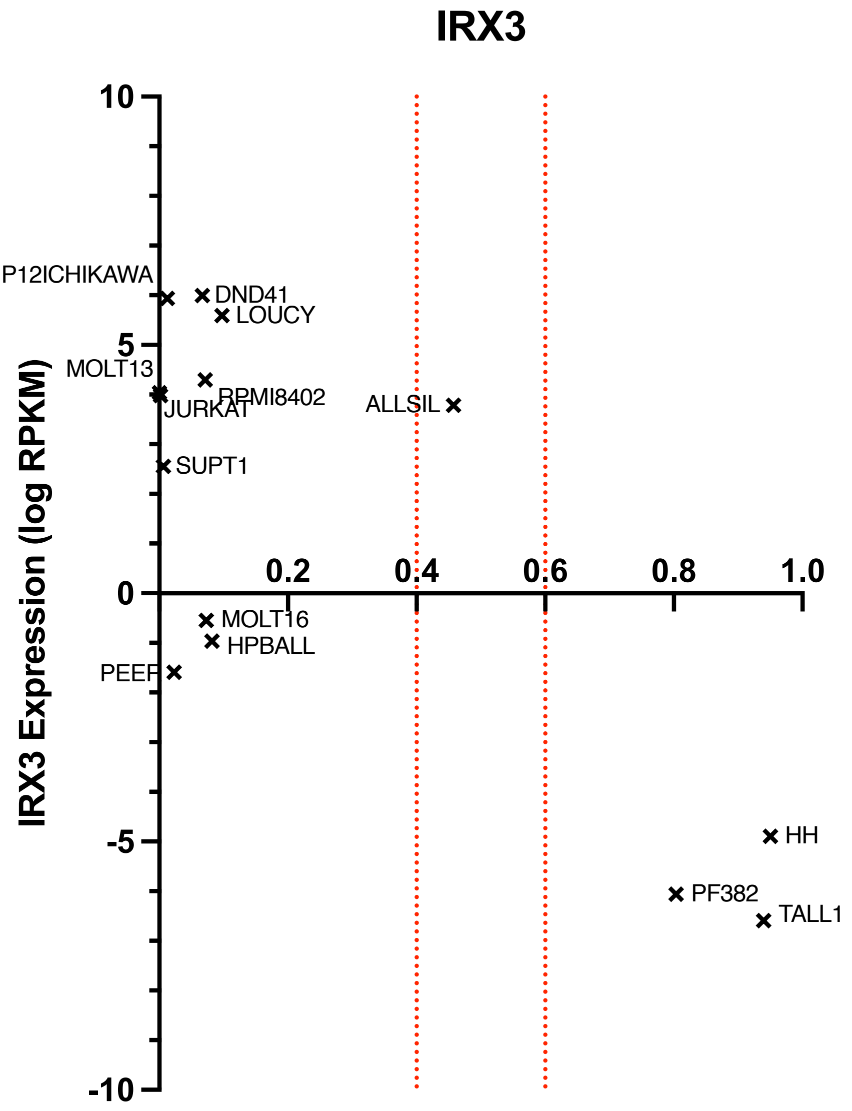


**Supplementary Figure 3. *IRX3* proximal promoter methylation from T-ALL cell lines against *IRX3* expression by RNA-Seq from Cancer Cell Line Encyclopedia.**
X-axis values refer to average promoter methylation within a 1 kb window upstream of the transcriptional start site, whereby a value of 0 is fully unmethylated and a value of 1 is fully methylated. Dotted red lines represent allelic imbalance range for promoter methylation. All promoter methylation values are averaged across maternal and paternal alleles for each sample. Y-axis values refer to *IRX3* expression (log RPKM) for T-ALL cell lines. This analysis assumes the *IRX3* locus is not subject to copy number alterations. Thus, absolute copy number calls were examined for 13 cell lines for which data was available in CCLE. HPB-ALL has a copy number of 1 for *IRX3* whilst all other cell lines had no copy number aberrations of *IRX3*. Note copy number data was unavailable for the TALL1 cell line in CCLE.

chr16:54050926-54051223 (+)
ACAGACAGTATGAAAACATTCTATGTGAAAGTTT**CCAACTGAAATC**GCACCACAAGCAGTTGGTTATCTGGA
GATAGACATCTTCAATTTCCTGATTGCATAATAGAAATGGGATTTCTTTTTCCTCTACAAAAAGCGTTATCCAT
AATGTTGACATAAAGTTTTCTGTGCCTTGAGTTCCAAAGAGTACTCTTCTCAAGAACAGCCCAGCATCTTCTCA
ACTGCTAACGGTTACTGCCATTTCTCCTCCTTCAAGGGAAAGGGTAAATGTTGCTAACATGATGAAGGCCAG
AAGAAG
**MYB_HUMAN.H11MO.0.A**
**ddPCR amplicon**

chr16: 54062930-54063133 (+)
TGCAACCCCAGTGACACAGGCAGCTGGAGCAAAGTGTGTGTGTGCACAC**ATGCGCCCCCGAGGGGT**GTGCG
GGGAGGTTAACTTCAGAGCCTTGTTTCCGAGAGGGTGTGTGTGGTGGTGCAGGTGGGATGAGAAGATTTCG
ACACTTTTCCTTTATGAAGTAGAACTGTGGATGTTGTGTCTTTTTGTTGGTAAGTTCAGGCA
**CTCF_HUMAN.H11MO.0.A**
ddPCR amplicon

**Supplementary Figure 4. Genomic locations of ddPCR probes relative to CTCF and MYB binding sites within *FTO* intron 8**
Coordinates positions are provided in hg38. Motifs were called by the Tomtom motif comparison tool from MEME suite using the HOCOMOCO Human (v11 CORE) library.

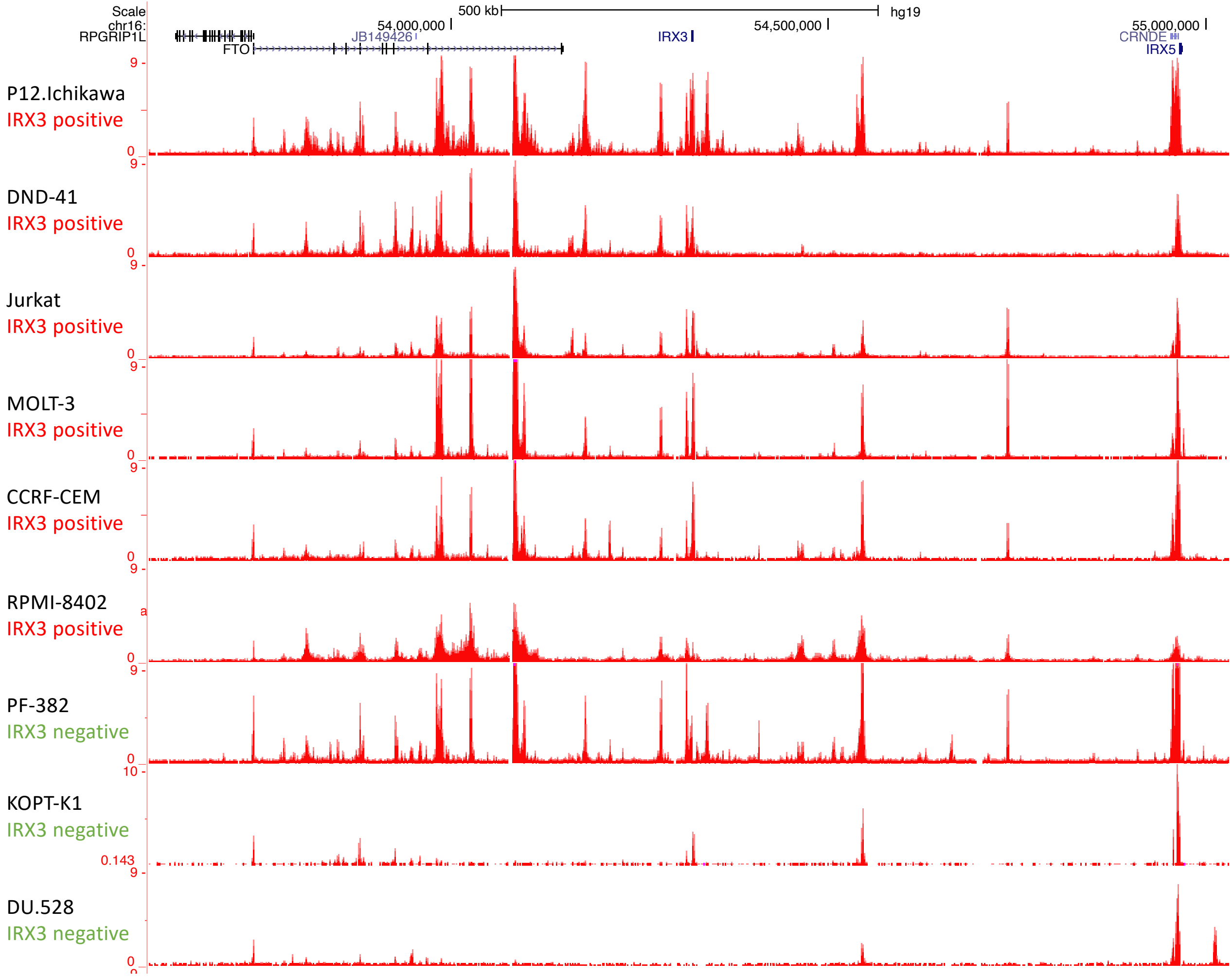


**Supplementary Figure 5. Comparison of enhancer architecture between an IRX3 positive and IRX3 negative T-ALL cell lines by ChIP-Seq**

H3K27ac ChIP-Seq data for the IRX3 positive T-ALL cell lines P12-Ichikawa, DND-41, Jurkat, MOLT-3, CCRF-CEM and RPMI-8402; and the IRX3 negative T-ALL cell lines PF382, KOPT-K1, and DU.528. Cell lines with similar enhancer positioning across the FTO/IRX3/CRNDE/IRX5 locus are highlighted by a blue bar (left) which includes the IRX3 negative cell line PF382. NCBI GEO accession #GSE76783. Genomic coordinates are in hg19.

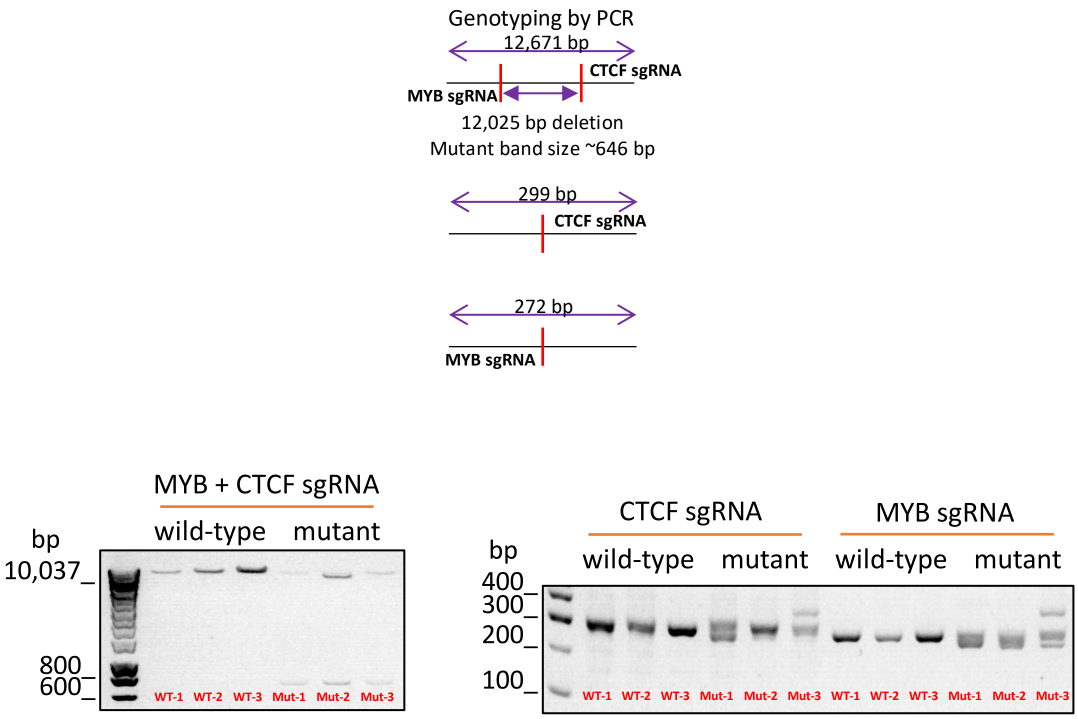


**Supplementary Figure 6. Genotyping of *FTO* intron 8 CRISPR/Cas9 edited single cell sorted clones**

(Top) Schematic outlining flanking PCR approach to identify single cell sorted clones with MYB and CTCF binding site deletions at *FTO* intron 8, and indels disrupting MYB or CTCF binding sites at *FTO* intron 8. This schematic is not to scale.

(Bottom) Agarose gel showing PCR products following PCR of gDNA extracted from single cell sorted clones. Deletion bands are observed ~646 bp for the MYB and CTCF deleted sites. Indel bands are observed for MYB or CTCF site disrupted clones. All single cell sorted clones derive from the PF-382 T-ALL cell line.

| Cell Line | *FTO* intron 8 site | Copy Number |
|---|---|---|
| P12 Ichikawa | CTCF | 2.03 |
| | MYB | 2.14 |
| CUTLL1 | CTCF | 2.52 |
| | MYB | 3.07 |
| DND41 | CTCF | 2.31 |
| | MYB | 1.91 |
| JURKAT | CTCF | 2.07 |
| | MYB | 2.15 |
| CTV1 | CTCF | 1.91 |
| | MYB | 1.78 |
| ALLSIL | CTCF | 0.97 |
| | MYB | 1.07 |
| PF382 | CTCF | 2.62 |
| | MYB | 2.56 |
| MOLT4 | CTCF | 1.93 |
| | MYB | 2.05 |
| MOLT3 | CTCF | 2.13 |
| | MYB | 2.16 |
| CCRFCEM | CTCF | 2.09 |
| | MYB | 2.18 |
| KARPAS45 | CTCF | 2.00 |
| | MYB | 2.19 |
| HSB2 | CTCF | 1.77 |
| | MYB | 1.88 |
| RPMI8402 | CTCF | 2.06 |
| | MYB | 2.19 |
| HPBALL | CTCF | 0.95 |
| | MYB | 0.98 |
| PEER | CTCF | 1.88 |
| | MYB | 1.91 |
| BE13 | CTCF | 2.05 |
| | MYB | 2.04 |
| KOPTK1 | CTCF | 1.72 |
| | MYB | 1.71 |
| DU528 | CTCF | 1.97 |
| | MYB | 1.98 |
| SUPT11 | CTCF | 2.01 |
| | MYB | 2.05 |
| LOUCY | CTCF | 2.04 |
| | MYB | 2.15 |
| SUPT1 | CTCF | 1.89 |
| | MYB | 2.02 |
| MOLT16 | CTCF | 1.91 |
| | MYB | 1.92 |

**Supplementary Table 1. Copy number calls at the CTCF and MYB binding sites within the *FTO* intron 8 locus by ddPCR in T-ALL cell lines**

Copy number calls of approximately 1 are highlighted in red. The genomic co-ordinates for the ddPCR probes are described in Supplementary Figure 3.

| gene | refseq | chromosome | start | aachange | class |
|---|---|---|---|---|---|
| **CTCF** | NM_006565 | chr16 | 67650782 | V363I | missense |
| **CTCF** | NM_006565 | chr16 | 67650762 | C356F | missense |
| **CTCF** | NM_006565 | chr16 | 67650767 | GPinsY358 | proteinIns |
| **CTCF** | NM_006565 | chr16 | 67645346 | T204fs | frameshift |
| **CTCF** | NM_006565 | chr16 | 67654639 | E376fs | frameshift |
| **CTCF** | NM_006565 | chr16 | 67645961 | H297fs | frameshift |
| **CTCF** | NM_006565 | chr16 | 67645346 | T204fs | frameshift |
| **CTCF** | NM_006565 | chr16 | 67654702 | H397Y | missense |
| **CTCF** | NM_006565 | chr16 | 67646024 | G318fs | frameshift |
| **CTCF** | NM_006565 | chr16 | 67655477 | A447fs | frameshift |
| **CTCF** | NM_006565 | chr16 | 67645399 | D222fs | frameshift |
| **CTCF** | NM_006565 | chr16 | 67645346 | T204fs | frameshift |
| **CTCF** | NM_006565 | chr16 | 67670634 | E627* | nonsense |

**Supplementary Table 2. CTCF mutations identified in primary T-ALL samples at diagnosis from the St Jude's cohort**
